# Detecting Horizontal Gene Transfer by Mapping Sequencing Reads Across Species Boundaries

**DOI:** 10.1101/039495

**Authors:** Kathrin Trappe, Tobias Marschall, Bernhard Y. Renard

## Abstract

Horizontal gene transfer (HGT) is a fundamental mechanism that enables organisms such as bacteria to directly transfer genetic material between distant species. This way, bacteria can acquire new traits such as antibiotic resistance or pathogenic toxins. Current bioinformatics approaches focus on the detection of past HGT events by exploring phylogenetic trees or genome composition inconsistencies. However, this normally requires the availability of finished and fully annotated genomes and of sufficiently large deviations that allow detection. Thus, these techniques are not widely applicable. Especially in an outbreak scenario where new HGT mediated pathogens emerge, there is need for fast and precise HGT detection. Next-generation sequencing (NGS) technologies can facilitate swift analysis of unknown pathogens but, to the best of our knowledge, so far no approach uses NGS data directly to detect HGTs.

We present Daisy, a novel mapping-based tool for HGT detection directly from NGS data. Daisy determines HGT boundaries with split-read mapping and evaluates candidate regions relying on read pair and coverage information. Daisy can successfully detect HGT regions with base pair resolution in both simulated and real data, and outperforms alternative approaches using a genome assembly of the reads. We see our approach as a powerful complement for a comprehensive analysis of HGT in the context of NGS data. Daisy is freely available from http://github.com/ktrappe/daisy.

## 1 Introduction

In bacteria, genetic material is commonly exchanged between organisms, a process known as *horizontal gene transfer* (HGT) or lateral gene transfer (Ochman et al., 2000, Boto, 2009, Wiedenbeck and Cohan, 2011). In contrast to vertical gene transfer, i.e. from one generation to the next, HGT enables the exchange of genetic material even between distant species mediated usually by transduction, transformation, or conjugation (Gyles and Boerlin, 2013). Via transduction or conjugation, the foreign DNA is carried in a plasmid or a bacteriophage, respectively, whereas via transformation, the recipient takes up nascent DNA from the environment. By means of HGT, complete genes and functional units, called insertion sequences (IS) or genomic islands (GIs), can be incorporated into the recipients’ genome. Each bacterium can also carry several phages at distinct phage insertion sites. Phages of the same type, e.g. λ phages, can also carry diverse genes in their replaceable region with the result that one bacterium can have multiple highly similar phages but with different gene content.

Not surprisingly, HGT greatly contributes to bacteria’s ability to adapt to changing environments (Hu et al., 2011, McElroy et al., 2014, Gyles and Boerlin, 2013). It has been demonstrated to play a major role for the acquisition of resistance to antibiotics (Barlow, 2009, Warnes et al., 2012). Moreover, HGT is not limited to bacteria but can also occur in vertebrates, including primates (and humans) (Crisp et al., 2015). However, the focus of the bioinformatics community with respect to HGT has mainly been on methods for detecting past HGT events (Ravenhall et al., 2015) from phylogenetic trees (e.g. Boc et al., 2010, Bansal et al., 2012) or based on genome composition (e.g. Metzler and Kalinina (2014), Jaron et al. (2013)). Composition properties such as GC content or k-mer frequencies usually deviate between different organisms and can therefore be used to detect sequence content of foreign origin. However, over time the foreign sequence signature ameliorates to its new host. Alien Hunter (Vernikos and Parkhill, 2006), e.g., therefore combines various compositional characteristics or *motiƒs* in a variable fashion, called Interpolated Variable Order Motifs (IVOM), to improve sensitivity. Their IVOM approach does not require gene annotation or gene position information and can hence be applied to newly sequenced genomes.

Common methods aim to retrace evolutionary history of finished bacterial genomes and have mostly been developed before next-generation sequencing (NGS) became available, and hence, do not directly use NGS data. NGS technologies are well established and widely used by now, and enormous amounts of NGS data is available in public repositories. NGS also offers the chance to detect HGT events early in analyses which can be important in outbreak scenarios. One prominent example is the EHEC outbreak in Germany back in 2011 (Frank et al., 2011), where non-pathogenic *Escherichia coli* bacteria residing in the gut of every human suddenly acquired two new toxins from another bacterium leading to excessive and often dangerous hemorrhagic gut infections. Especially here, identification and characterization of the pathogen causing the outbreak is highly important. Fast and reliable pathogen identification or detection of antibiotic resistance are generally of particular interest (Byrd et al., 2014). Important applications in diagnostics in the context of HGT are the detection of novel bacterial strains evolved through HGT or the distinction of a single infection with such a strain from a parallel infection by two different strains (Fricke and Rasko, 2013). This distinction is important for treatment and to prevent spreading of the disease, especially with the more frequently occurring cases of antibiotic resistances. With a special focus on these applications, we developed an HGT detection tool that directly uses NGS data.

While methods that directly address the detection of HGT events from NGS data are lacking, various methods for finding structural variations (SVs) in human exist, as for instance reviewed by Medvedev et al. (2009), Alkan et al. (2011), and Pabinger et al. (2014). Furthermore, first systematic attempts are being made to transfer methods for SV discovery to other species, including plants (Leung et al., 2015) and bacteria (Barrick et al., 2014, Hawkey et al., 2015). The latter approaches focus on detecting SVs *within* a genome and do not aim to detect the transfer of genetic material *between* species. To our knowledge, no such method exists to date (Ravenhall et al., 2015).

Conceptually, detecting an HGT event has similarities to identifying an inter-chromosomal translocation in an organism with multiple chromosomes (such as human). Nonetheless, a number of differences render existing methods not directly applicable for the purpose of detecting HGT events. On the one hand, the underlying mechanisms are different, e.g. phage-mediated transfers versus integration of nascent DNA, which potentially leads to other breakpoint signatures. On the other hand, bacteria are subject to much higher mutation rates than humans and can undergo faster evolution (Lee et al., 2012). This usually also implies fast divergence of sequences acquired via HGT (Iranzo et al., 2014). Besides sequence deviation due to evolution, reference databases still contain a number of draft genomes or mis-assemblies (Salzberg and Yorke, 2005, Kuhring et al., 2015), adding sequence deviation due to technical artifacts. Usually, the peformance of methods for calling structural variations from human NGS data deteriorates in the presence of large amounts of sequence divergence. Despite these issues, structural variant detection methods, which we briefly survey below, provide an excellent starting point to approach HGT detection when we combine their individual strengths.

The most commonly used methods to detect structural variants from NGS reads are based on mapping reads to reference genomes. To this end, three different paradigms exist: (i) *Coverage* information can be used to detect copy number variants (e.g. Abyzov et al. (2011), Miller et al. (2011)). This allows finding regions that are covered by significantly more or less reads than the background genome in order to predict copy number gains or losses. Such approaches are effective for large events, usually starting from approximately 5 kb, and work best if multiple samples are available for comparison, allowing for properly handling coverage biases (Dohm et al., 2008). (ii) The second class of approaches leverages *read pair* information. Here, the idea is to detect deviation from the expected relative mapping positions of two paired reads generated by mate pair or paired-end sequencing. This technique allows for uncovering also copy neutral events such as inversions, or copy-neutral translocations. The accuracy in terms of breakpoint placement and event length strongly depends on the insert size distribution of the library. In practice, approaches that first classify read pairs as concordant or discordant and then make predictions based on the discordant reads (e.g. Chen et al. (2009), Hormozdiari et al. (2010)) are usually effective for events of approximately 250 bp and larger, while approaches that use all reads (e.g. Lee et al. (2009), Marschall et al. (2012)) can predict variants starting from approximately 30 bp. (iii) Finally, it is possible to align reads across SV breakpoints, which is often referred to as *split alignment* or *split-read mapping* (Trappe et al., 2014, Emde et al., 2012, Ye et al., 2009, Marschall and Schönhuth, 2013, Karakoc et al., 2012). Such approaches can deliver single base pair resolution, but have limitations with respect to repetitive regions: Splitting the reads makes alignment ambiguity even more likely to occur than for full length reads. Especially split-read approaches then have to trade sensitivity for high numbers of false positive calls.

The different paradigms outlined above have different strengths and weaknesses and use different information sources. Therefore, many *hybrid* techniques that use more than one of these ideas have been developed in the past years (e.g. Rausch et al. (2012), Marschall et al. (2013), Jiang et al. (2012)). In contrast to these hybrid approaches, that integrate different techniques into one algorithm, *meta tools* provide a unifying platform to integrate the results of complementary methods into a unified variant call set (Lin et al., 2014, Leung et al., 2015).

Besides mapping reads to reference genomes for SV detection, it is also possible to subject them to *de novo* assembly (Luo et al., 2012a, Bankevich et al., 2012, Zerbino and Birney, 2008). There are many advantages to this approach, including that biases due to the choice of reference are avoided, all classes of SVs can be addressed, and 1 bp resolution is attained. However, these advantages only apply *iƒ* the reads can be assembled into sufficiently long contigs, which cannot always be achieved from short read data. Although long read sequencing technologies can drastically improve the ability to assemble difficult regions (Chaisson et al., 2015), short read technologies are still more prevalent, more cost effective and will thus continue to play a major role in the coming years. This holds in particular for the application to fast evolving genomes such as bacteria since here the low technical error rates of short reads is of clear advantage. Hence, short read technologies are most likely the method of choice in outbreak situations.

### Contributions

In this manuscript, we introduce Daisy, a novel mapping-based HGT detection tool using NGS data. Daisy facilitates HGT detection in outbreak scenarios such as the EHEC outbreak 2011 in Germany. Outbreak situations require fast and reliable characterization of novel or unknown pathogens, or the distinction of such a novel pathogen from double infections to prevent disease spreading and to apply proper treatment. We incorporate all three paradigms of mapping-based SV detection: We identify HGT boundaries with split-read mapping and then filter candidate regions using coverage and read pair information (see Figure 1). The identification part ensures sensitivity in the presence of sequence divergence whereas the filtering part removes un-specific, non-HGT related events. We show the utility of mapping-based techniques for HGT detection by applying our approach to one simulated data set and two different bacterial case studies, each showing that mapping can help beyond what can be achieved with assembly for HGT detection. With Daisy, we provide an easy to use open source software relying on community standards such as VCF files and readily usable output.

**Figure 1:**
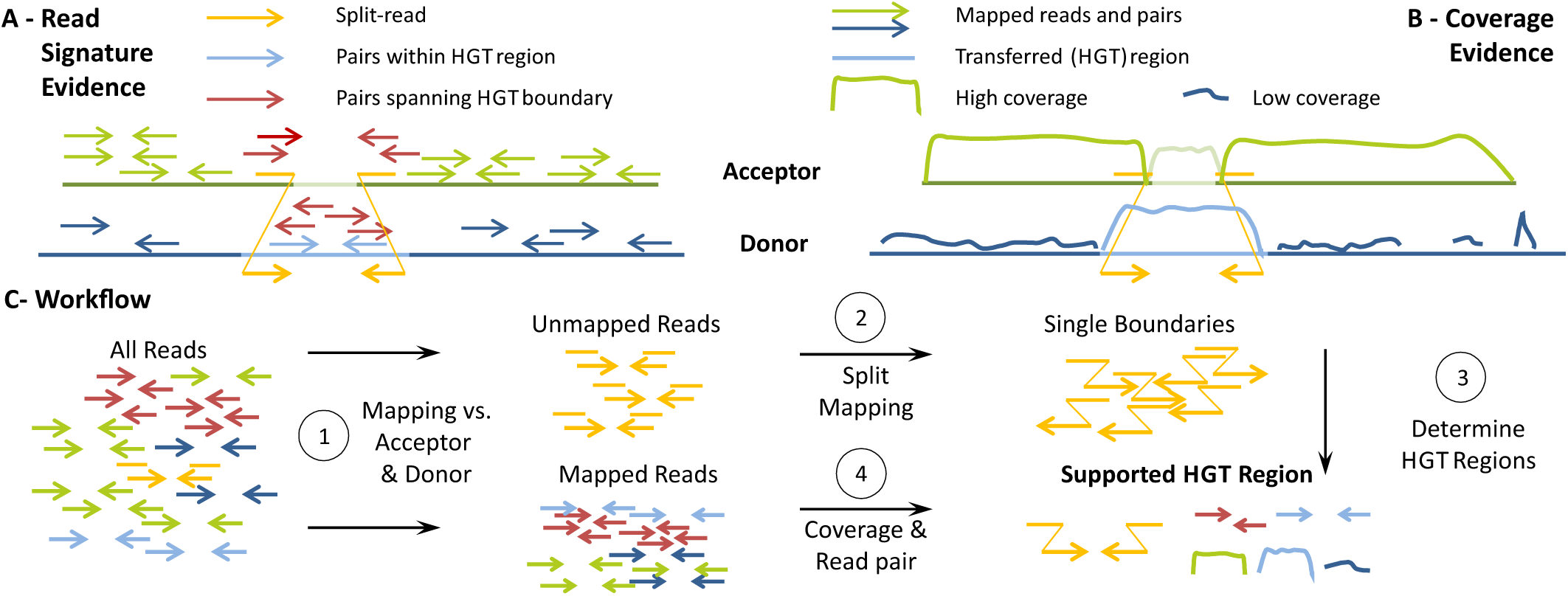
Daisy evidence and workflow. (A) Mapping evidence based on read signature: For an acceptor genome (green) and a donor genome (dark blue) with the transferred region (light blue), we evaluate read mapping information from split-reads (yellow arrows) crossing the transfer boundaries, pairs exclusively mapping within the transferred region (light blue arrows), and read pairs spanning the boundary, i.e. they have one read on either side of the boundary (dark red arrows). (B) Mapping evidence based on coverage: We evaluate the coverage based on acceptor reads (green arrows) and donor reads (dark blue arrows). We expect the coverage of the acceptor genome to be high and homogeneous (green lines) except for the HGT insertion site (light green line). The coverage in the transferred region (light blue line) should be comparable to the acceptor genome and higher than the coverage in the remaining donor region (dark blue lines). (C) Workflow overview: After initial read mapping (1), unmapped reads are split mapped to determine single HGT boundaries (2). Single boundaries are paired up to form candidate regions according to size constrains (3), and evaluated in accordance with mapping information regarding coverage and read pair signatures (4).

## 2 Methods

Daisy is a comprehensive, mapping-based tool for HGT detection using sequencing data of an HGT organism, i.e. an organism with an acquired HGT, without a complete reference sequence available. The input is a set of sequencing reads from an HGT organism, i.e. an organism with a suspected, but not yet known or identified HGT event. With no complete reference available that includes the HGT event, we map the reads against the acceptor genome reference (the parent genome of the HGT organism acquiring the HGT sequence) and the donor genome reference (the parent donating the HGT sequence; see Figure 1).

We assume that donor and acceptor references are known. In a first read mapping step, Daisy identifies possible split-read candidates and also acquires mapping information around the HGT boundaries that is later in-corporated for HGT support (step 1 of the Workflow in Figure 1C, details below). We use a dedicated split-read mapper to determine the single boundaries (step 2), then pair up the boundaries to HGT regions according to size constraints (step 3), and integrate the mapping information of read pairs spanning and mapping within the HGT region (step 4). We further filter the results by a bootstrap based approach where we resample coverage and the number of reads spanning or within the HGT boundaries from random regions in acceptor and donor. As an additional step, we map the read pairs of candidate donor regions against a bacteriophage database, and flag those candidates having relevant hits. All candidate regions meeting a pre-defined support threshold are reported in VCF format. Daisy has two modes. The automated mode supports single acceptor and donor references with full filtering options. The manual mode gives the possibility to examine multiple donor genomes (see data set KO11FL for an example), although without filtering.

### Read filtering

We simultaneously map the reads against the acceptor and the donor genome to identify the fully mapping reads. We use Yara (successor of Masai, Siragusa et al. (2013)), a read mapper designed with focus on efficiency and low-error mapping. Hence, Yara does not incorporate partial mapping or bad-quality last resort mapping. As a result, Yara maps less of these reads compared to other mappers (data not shown) and more split-read candidates are investigated during split-read mapping.

### Split-read mapping

In an HGT event, a part of the donor genome has been integrated into the acceptor genome. Given a set of reads of the HGT organism, we expect to see reads mapping across the HGT boundary where one part of a read maps to the acceptor and the other part to the HGT origin in the donor (see yellow reads in Figure 1). When these reads are split-read mapped concurrently to both acceptor and donor, we can identify the breakpoints of an HGT event because the signature of an HGT in SV terms then resembles an inter-chromosomal translocation.

These HGT breakpoints are also the main evidence for an actual integration of the possible transferred region in contrast to potential contamination or co-existences of both donor and acceptor. We use the SV detection tool Gustaf (Trappe et al., 2014). Gustaf works with single-end data but also incorporates paired-end information from paired-end data. It can handle multiple splits per read and alignment gaps at the read ends or in the middle of the read which is an important property in view of the high bacterial evolutionary rate and common micro-homologies at breakpoint locations. The expected number of split-reads depends on the read coverage and the evolutionary distance between the sequenced organism and its putative acceptor and donor genomes used for analysis. The default value of the user definable parameter for the required number of split-reads is therefore set to 3 (very sensitive but avoiding random split-reads).

### Candidate identification

The single breakpoints from the split-read mapping give possible start and end positions, in both acceptor and donor, of an HGT event. The combination of these start and end positions is subject to size constrains regarding the regions delimited in the acceptor and donor genomes in order to sensibly restrict the number of candidate regions. Depending on whether only single genes, operons, or complete bachte-riophages are transferred, these regions can vary largely in size and range from a few hundred to several thousand base pairs. The delimited region in the acceptor genome can also be equally large as the designated HGT region if, e.g., another bacteriophage is occupying the destined phage insertion site there. The values for minimal and maximal HGT size are therefore parameterized and user definable. Default values used in the benchmarks are 500bp and 55,000bp for minimal and maximal HGT size, respectively. We also reduce duplicate entries. Once we identified a valid candidate, we remove any further identified candidates within a base pair range of a specified tolerance (default 20 bp) around acceptor and donor start and end positions.

### Coverage and read pair integration

Each candidate region is then examined for additional mapping support regarding mean coverage, number of pairs spanning and within HGT boundaries (see “Mapping Evidence” in Figure 1). Coverage can vary due to extreme GC content, sequencing efficiency or rearrangement events such as induced by a HGT.

Theoretically, the expected coverage of the acceptor genome should be equally high and homogeneous as the sequencing coverage of the HGT organism (depicted as “High coverage” in Figure 1), except for the HGT insertion site. The coverage in the HGT insertion site should be either much lower because the sequence content is unrelated to HGT organism and donor or much higher because it is occupied by another related phage. The coverage in the donor HGT region should, again theoretically, resemble the coverage of the acceptor whereas the coverage of the remaining donor should be low (depicted as “Low coverage” in Figure 1). Depending on the evolutionary distance between HGT organism, donor and acceptor, the observed coverage properties of the region can deviate. A direct statistical comparison, e.g. using the framework proposed in Lindner et al. (2013), may lead to insignificantly small values, even for the true HGT regions. We therefore introduced a bootstrap like resampling method where we test the candidate regions compared to equally sized random regions in acceptor and donor. The default sampling size is 100 random regions. As stated above, the donor region coverage should be higher than the coverage of the remaining donor. Per default, we require the donor region mean coverage to be higher than the coverage of the random donor regions in at least 95% of the cases, i.e. to have a bootstrap result of ≥ 95. Again, the acceptor region should be unrelated (low coverage) or also have phage origin (high coverage). Hence, we require the mean coverage to be either higher (alternative phage) or lower (unrelated sequence) than the random region coverages in at least 95%, i.e. the bootstrap value has to be (≥ 95 or ≤ 5).

In addition to coverage evidence, we also incorporate mapping evidence from read pairs that are spanning the HGT boundaries (dark red reads in Figure1) and those that map completely within the HGT boundaries of the donor (light blue reads). For the spanning pairs, one mate is mapping on one site of the boundary outside the HGT region in the acceptor whereas the other is mapping on the other site of the boundary inside the HGT region in the donor. Both have to map within a range of half the defined maximal HGT size from the boundary (i.e. a total range of the maximal HGT size around the boundary). For the pairs within, we compare the number of pairs within the boundaries to the number within equally sized random regions where we expect the HGT region to have more such pairs than the random regions. We apply the same idea of a resampling method from the coverage evidence (using the same random regions) for the evidence from mapped read pairs spanning and within the HGT boundaries. The required resampling value is also 95.

### Bacteriophage screening

If the HGT was phage mediated and HGT organism and acceptor contain, and maybe share, several similar phages, the results obtained via split-read mapping can be ambiguous (see Figure 2). After filtering the HGT candidates, we therefore screen the EBI phage database (Brooksbank et al. (2014)) from the European Nucleotide Archive (ENA) (Leinonen et al., 2011) for evidence of the candidates’ donor HGT regions. We first map all reads against the phage references and during the screening, we evaluate if the reads mapping within or across the donor HGT region also map to any database entry and report this percentage in the TSV output file (see below). This step is not a filter step but intended as an additional flag for each candidate.

The filtered candidates are written to a VCF output file (Danecek et al., 2011), all candidates with bootstrap information are written to a TSV file.

**Figure 2:**
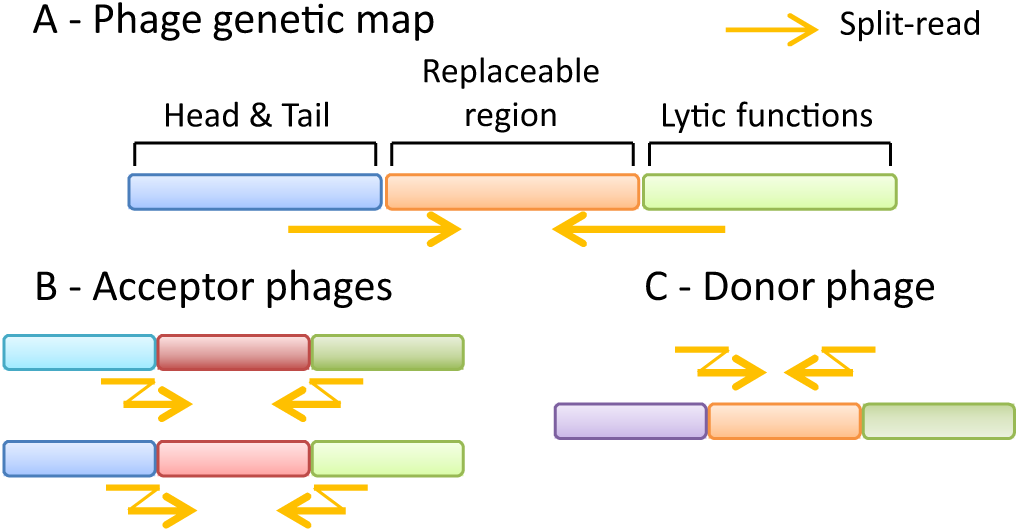
Phage composition. (A) Basic genetic map of a λ phage: Each λ phage has mosaic like coding regions for head, tail (blue), and lytic functions (green). In addition, the phage may contain an interchangeable region (orange). HGT related split-reads (yellow) cross the border between these regions. (B) The acceptor is likely to carry similar phages (variants of blue and green) with another replaceable content (red). This can make the split-read mapping ambiguous between the blue and green parts, respectively. (C) The donor very likely carries the transferred HGT region (orange) in a similar phage (blue and green) as the HGT organism.

## 3 Experimental Setup

### Datasets

We tested our method on one simulated and two real data sets, each containing an HGT event with distinct challenges.

#### H. pylori

The *Helicobacter pylori* data set is a simulated set. Here, we chose *Escherichia coli* K12 as the acceptor and *H. pylori* strain HPML01 (Wang et al., 2015) as the donor. We introduced SNPs, small and large indels using the simulator Mason2 (Holtgrewe, 2010, 2014) into *E. coli* K12 and into the phage-like sequence of *H. pylori*. We simulated 150bp paired-end reads with Illumina error profile and 100x coverage with Mason2.

#### KO11FL

The first real data set includes *E. coli* W as the acceptor and *Zymomonas mobilis* as the donor genome as well as the cloning vector pBEN77 as a second donor, the resulting genome is the transgenic *E. coli* KO11FL (Turner et al., 2012). The KO11FL is a laboratory version of the original transgenic KO11 (Ohta et al., 1991). *E. coli* W is the parent strain of KO11FL which contains a cloned operon *pcl* including the genes *pdc* and *adhB* from *Z. mobilis*, and a *cat* gene not present in *Z. mobilis*. We therefore chose pBEN77 as a donor genome for the *cat* gene. This transgenic biotechnology scenario resembles a natural HGT event and gives the necessary ground truth on real data.

Figure 3 depicts the composition of *E. coli* KO11FL with the transferred genes *pdc* (green), *adhB* (red) and *cat*, the target site of the acceptor genome *E. coli* W (insertion breakpoint framed purple), and excerpts of the donor genomes *Z. mobilis* and cloning vector pBEN77 indicating the positions of the transferred genes. The purple framed HGT region in *E. coli* KO11FL has 20 consecutive copies. The exact order, orientation and positions of the segments enumerated with I-IX has been determined with BLAST (megablast, default parameters). The adjacency of segments I and II defines the first HGT boundary and the adjacency of V and VI defines the second boundary number. The important and challenging part is that II and VI belong to two different donors, i.e. we have a transition within the HGT region and cannot define a single candidate region by pairing up single boundaries as we did for the *H. pylori* data. However, since the transfer was very recent and the boundaries are still clear enough, we will aim to detect all HGT related boundaries via split-read mapping alone.

**Figure 3:**
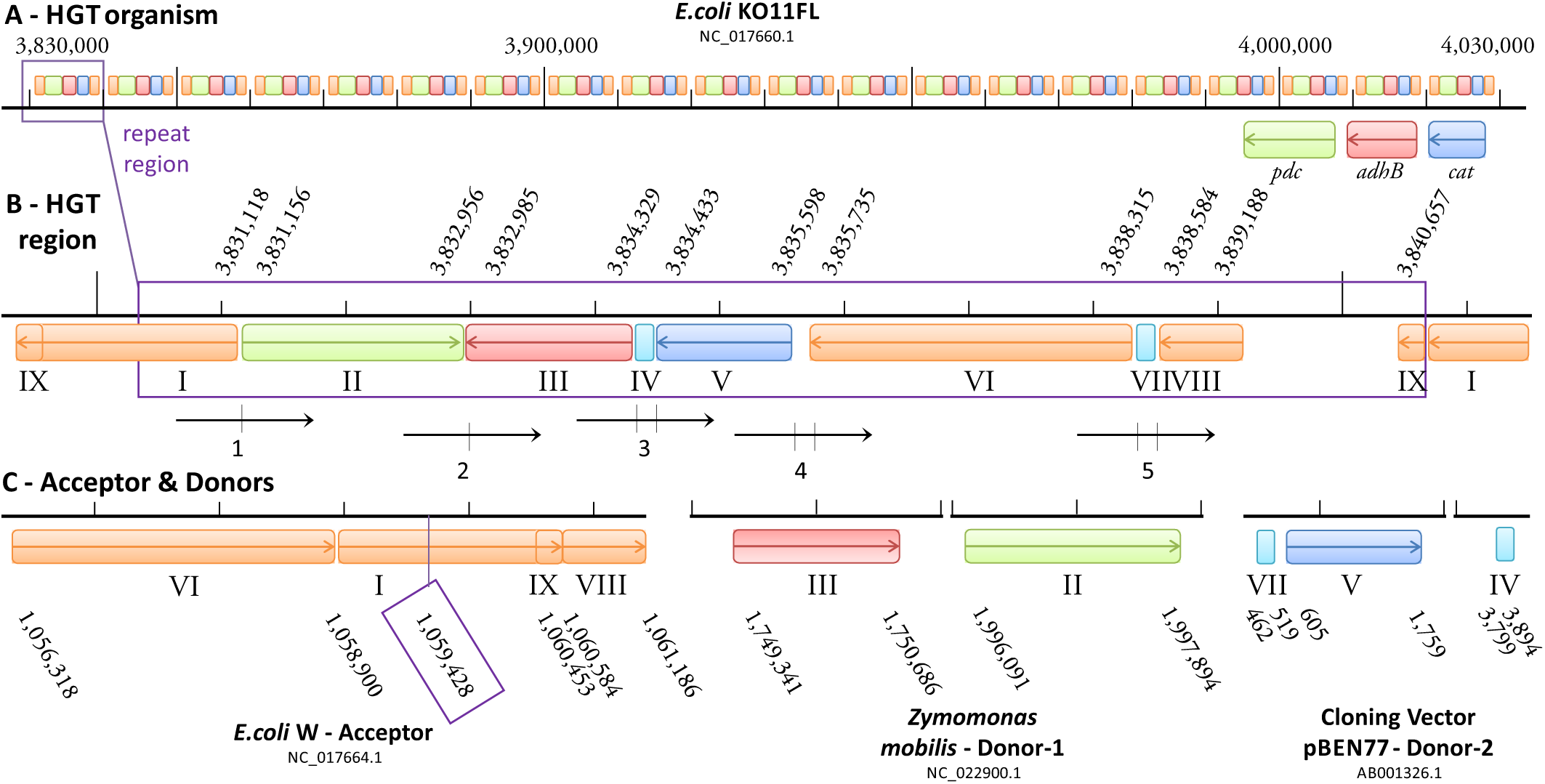
KO11FL composition of HGT region. (A) HGT organism *E. coli* KO11FL: The transgenic KO11FL has 20 copies of the transferred HGT region enclosed by the purple rectangle. (B) HGT region composition of KO11FL: Shown are positions of the transgenic genes *pdc* (green), *adhB* (red) and *cat* (blue) and adjacent segments (I-IX) within *E. coli* KO11FL. Reads enumerated 1-5 span HGT related adjacencies of segments I-IX, two dashes on a read imply multiple adjacencies or gaps, resulting in multiple splits of the read. (C) Acceptor & donor HGT region composition: Shown are positions of *pdc, adhB*, and *cat* in the donor references *Z. mobilis* and pBEN77, and the HGT insertion site in acceptor reference *E. coli* W. Note the different order and orientation of some of the segments I-IX compared to (B). All positions in (A)-(C) were determined with BLAST.

The read types numbered 1-5 in Figure3 are the expected split reads relevant for or related to HGT detection. Reads 3-5 have multiple splits indicated by two dashes, i.e. the split read mapper must be able to handle multiple split reads. For read 4, the middle part of the read, 137 bp, is covered by neither acceptor nor donor genomes, i.e. the split-read mapper has to also handle such scenarios. Reads 3-5 are multiply split and read 4 spans a gap of over 130 bp, which shows the necessity of a sensitive and versatile split-read mapper. Adjacencies II-III and IV-V reflect intra-chromosomal rearrangements in *Z. mobilis* (II-III) and pBEN77 (IV-V) and are not part of this evaluation. KO11FL has been originally assembled from Roche 454 reads (Turner et al., 2012). Contig gaps have been filled by PCR and Sanger sequencing, and then resequenced Illumina short read paired-end data has been assembled using the Roche 454 assembly as a template. However, only the Roche 454 reads are available via the SRA (SRX022824) and have been used in our benchmark.

#### EHEC

The second real data set comprises the *E. coli* O157:H7 Sakai strain as an HGT organism. The *E. coli* O157:H7 serotype is associated with diseases most often and the Sakai strain has been sequenced from an outbreak in Japan (Zhang et al., 2007). *E. coli* O157:H7 arose from the enteropathogenic *E. coli* O55:H7, the acceptor, acquiring Shiga-Toxins (Stx) via HGT of a lam-doid phage in the sequential evolution from its progenitor (Kyle et al., 2012). *E. coli* strains that have both Stx1 and Stx2 have been shown to carry them in two separate and distinct (Herold et al., 2004) lambdoid phages (Allison et al., 2003). The *Shigella dysenteriae*, the assumed donor, is the only Shigella serotype carrying Stx (Yang, 2005). Stx1 is almost identical to the *Shigella dysenteriae* toxin (Shaikh and Tarr, 2003), whereas Stx2 only shares up to 60% with Stx1. Stx in *S. dysenteriae* at positions 1283705-1285203 is carried by the lambdoid stx-phage P27 (all Stx phages are lambdoid bacteriophages (Smith et al., 2012)).

In *E. coli* O157:H7 Sakai, the Stx1 phage Sp15 occupies the insertion site *yehV*, Stx2 phage Sp5 insertion site *wrbA* (Kyle et al., 2012). In *E. coli* O55:H7, *yehV* is occupied by another lambdoid phage (Cp10, see Table S1 in Kyle et al. (2012)), whereas *wrbA* is still intact (i.e., there is no phage at this insertion site, and the *wrbA* protein coding gene is intact (Shaikh and Tarr, 2003)).

### Assembly Approach

A comparable approach for HGT detection is de novo assembly with subsequent whole-genome analysis. We chose SOAPdenovo2 (Luo et al., 2012b) as a suitable assembler, in particular for short-read Illumina data (GAGE, Salzberg et al. (2012)). We assembled the reads of all data sets with parameters SOAPdenovo-127mer all-R-F-u-K 31-m 91.

To further evaluate the assembly results, we applied BWA-MEM (Li and Durbin, 2009) in order to detect possible HGT breakpoints directly on the scaffolds, and Alien Hunter (Vernikos and Parkhill, 2006) as an composition-based HGT detection tool to detect possible GIs. Since Alien Hunter is designed for fully sequenced and annotated genomes, we also applied Alien Hunter to the HGT organism reference genomes (i.e., the simulated *H. pylori* genome, *E. coli* O157:H7, and *E. coli* KO11FL) for evaluation purposes, although in our general use case, we assume that the HGT organism has not been sequenced yet or is unknown.

The detected regions from Alien Hunter have a score and are ranked in descending order. The higher the score, the more likely the region matches a foreign genomic island. Daisy and BWA-MEM candidate regions are not scored and therefore not ranked. For Daisy, we consider valid HGT candidates passing the 95% sampling threshold. For BWA-MEM, we consider all regions conforming the same size constraints as for Daisy.

## 4 Results

### H. pylori

Daisy finds one true positive (TP) HGT candidate with base pair precision without any false positives (FPs) (see Table 2 (A) *H. pylori*). Of the spanning read pairs and pairs within, 53% are mapping to the bacteriophage database. This indicates a phage-natured origin of the donor-region which conforms with the ground truth of the inserted Helicobacter phage 1961P-like sequence.

### Assembly

Assembly of the simulated reads resulted in 23,936 contigs, 6,484 of them covered by one of the 17,452 scaffolds (N50 of 89,444). BWA-MEM also only reports the correct breakpoints with base pair precision and without FPs. Alien Hunter detects the region on both the complete genome and the respective assembly scaffold but finds another 63 alternative hits on the assembly and 62 on the complete genome. The regions depicted by Alien Hunter deviate over 2,200 and up to 5,108 bp regarding the true start and end positions. All three tools detect the HGT region as the best (Alien Hunter) or only candidate but only Daisy and BWA-MEM with base pair precision.

### KO11FL

The transgenic *E. coli* KO11FL is a case study with a very recent artificial transfer and, due to the transgenic origin, two donor references. The composition of the HGT region is shown and explained in Figure3. The read types labeled 1-5 in Figure3 cover the HGT related boundaries which leaves adjacencies I-II, III-IV, V-VI, VI-VII, and VII-VIII as ground truth. We applied the manual mode of Daisy to investigate the HGT related boundaries.

A summary of the reported breakpoints is listed in Table 1 with references colored according to Figure 3 for easier reference. The five highest ranked TPs cover all adjacencies stated above, and, likely due to the 20 copies, have distinct high support of over 170 up to 3,294 reads whereas the FPs attain support values only up to 59, allowing perfect separation by a simple cutoff. Furthermore, most FPs can be assigned to an adjacency not expected based on the ground truth shown in Figure 3. We cannot assess whether some of these additional adjacencies reflect alternative compositions of the respective components in some of the 20 copies.

**Table 1:**
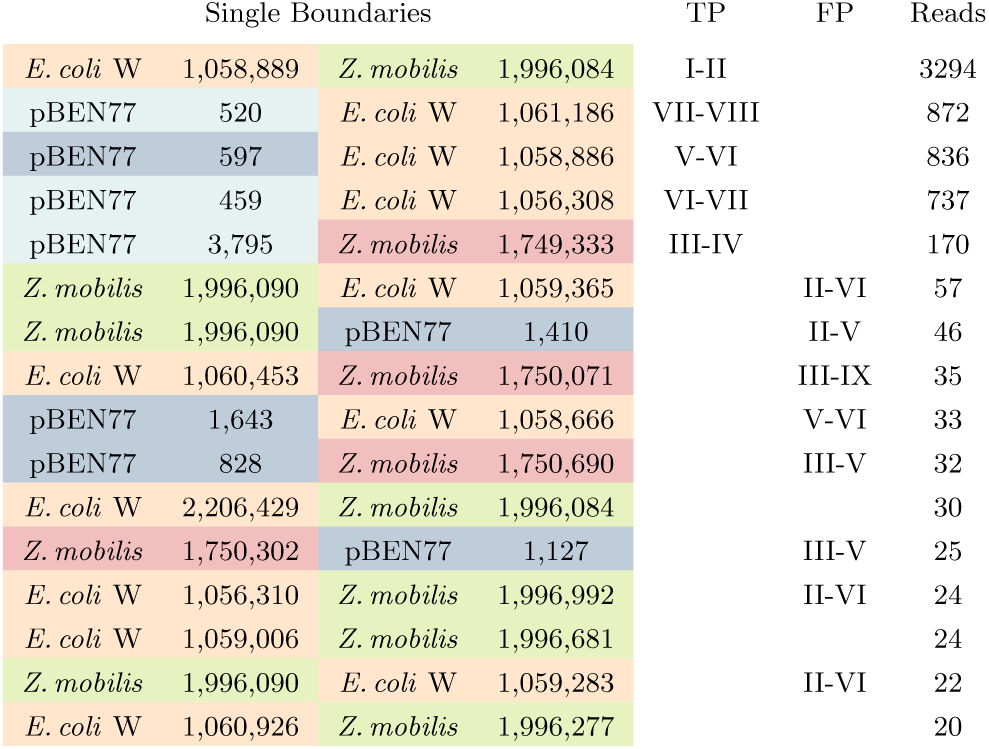
Daisys KO11FL results: References are colored according to Figure 3. Column TP (true positives) states which of the adjacencies between segments I-IX in Figure 3 the boundary covers, if any. Column FP (false positives) states possible adjacencies when considering alternative repeat region compositions within the 20 copies in *E. coli* KO11FL, empty entries are unrelated FPs. The column Reads states the number of split-reads supporting the translocation.

Table 2 (B) states the total number of breakpoints (16) as the *number oƒ hits* with a total of five true positives. As breakpoint distances on donor and acceptor, we calculated the mean distance of all true positive breakpoints involving acceptor-donor boundaries (adjacencies I-II and V-VI) since these enclose the region and can therefore be compared with Alien Hunter.

**Table 2:** Results for Daisy compared to Alien Hunter and BWA-MEM: For evaluation purposes, Alien Hunter has been applied to both the assembly and the full reference genome. Column *true region detected* states if the method was able to find the correct HGT region. Column *number of alternative hits* reports the total number of candidates detected. For the KO11FL, we additionally state the number of true positives (TP). In the last three columns, we state the precision of the correct candidate in terms of breakpoint distance. Note that Alien Hunter reports candidate regions with regard to the reference HGT organism whereas Daisy and BWA-MEM report breakpoints on acceptor and donor. For Alien Hunter, we calculate the base pair distance of the correct candidate region to the ground truth on the HGT organisms reference (column *distance true region*), for Daisy and BWA-MEM, we calculate the base pair distance on acceptor and donor (columns *breakpoint distance acceptor and donor*) (in the form start distance/end distance). Due to the ambiguous positioning of the HGT genes within the phage(s), columns with * state zero distance if breakpoints lie within phage region. (A) For the simulated *H. pylori* data set, all three methods are able to detect the HGT region as the best (Alien Hunter) or only candidate but only Daisy and BWA-MEM with base pair precision. (B) For KO11FL, an assembly using SOAPdenovo2 did not produce any scaffolds, and Alien Hunter was applied to the KO11FL genome. Both Daisy and Alien Hunter then detect the region. (C) For the EHEC data set, finds the true candidate. Alien Hunter finds 382 candidate regions when applied to the assembly but non is matching the scaffold with the HGT region. With BWA-MEM, we find 177 candidate regions, the closest is overlapping the true HGT region but without breakpoint precision.

| Tool & Data | True Region Detected | Number Hits | Distance to True Region (start/end) | Breakpoint Distance Acceptor (start/end) | Breakpoint Distance Donor (start/end) |
| --- | --- | --- | --- | --- | --- |
| **(A) *H. pylori*** |  |  |  |  |  |
| Daisy (reads) | yes | 1 | n/a | 0/0 | 0/0 |
| Alien_Hunter (genome) | yes | 63 | 2,289/2,215 | n/a | n/a |
| Alien_Hunter (assembly) | yes | 64 | 5,108/2,888 | n/a | n/a |
| BWA-MEM (assembly) | yes | 1 | n/a | 0/0 | 0/0 |
| **(B) KO11FL** |  |  |  |  |  |
| Daisy (reads) | yes | 16 (5 TP) | n/a | 39/14 | 22/8 |
| Alien_Hunter (genome) | yes | 109 (15 TP) | 1,732/1,174 | n/a | n/a |
| Alien_Hunter (assembly) |  |  | assembly failed |  |  |
| BWA-MEM (assembly) |  |  | assembly failed |  |  |
| **(C) EHEC** |  |  |  |  |  |
| Daisy (reads) | yes | 6 | n/a | 0/0 * | 32/2,876 |
| Alien_Hunter (genome) | yes (Stx2) | 93 | 0/0 * | n/a | n/a |
|  | yes (Stx1) | 93 | 0/0 * | n/a | n/a |
| Alien_Hunter (assembly) | no | 382 | — | — | — |
| BWA-MEM (assembly) | yes | 177 | n/a | 0/0 * | 1,864/39,049 |

We could successfully detect all of the five possible split-read types including the multiple split ones (reads 3-5 in Figure3) and the one read type covering a gap of over 130 bp (read 4). However, we could not verify the results with the HGT filter due to the two-donor scenario. Still, this case study shows that, given sequencing data of recent HGTs, it is possible to determine the right boundaries with accurate base pair resolution and high confidence even by split-read mapping alone.

#### Assembly

Assembly of the 454 reads from the KO11FL data set with SOAPdenovo2 resulted in 455,419 contigs (singletons) of up to 1,011 bp. No scaffold was constructed, likely due to the repetitive nature of the genome. Turner et al. (2012) also pointed out the long gaps between contigs in their assembly of this data set which they had to fill with PCR and additional Sanger sequencing. Due to the failed assembly, we did not apply Alien Hunter or BWA-MEM to the contigs.

When applied to the finished full KO11FL genome instead of the assembly, Alien Hunter finds a total of 109 potential GIs where 15 of them overlap with 15 of the 20 copies of the transferred genes *pdc, adhB*, and *cat* in KO11FL (see also Figure 3 A - HGT organism and B-HGT region). The mean distance for start and end position of this 4,442bp region, however, is 1,732 (start) and 1,174 (end). This is much higher than the mean distances for Daisy. Since we assume that the full genome is not available, Daisy outperforms the assembly approach even for the longer 454 single-end reads.

### EHEC

For the EHEC data set, the true transfer according to literature (Kyle et al., 2012) and BLAST hit examination is 2,643,556-2,694,691 in *E. coli* O55:H7, where the λ phage Cp10 occupies the phage insertion site *yehV*, to positions 1,283,705-1,285,203 in *S. dysenteriae* Sd197 where a defective prophage carries the Stx genes.

Applying Daisy, we created 145 candidates of which six passed the resampling filter. The true acceptor positions from *E. coli* O55:H7 stated above are not among them, and only one pair of donor positions matches the stated *S. dysenteriae* positions. The true acceptor positions are among the remaining filtered out candidates but have very low bootstrap values. So at first glance, it seems as if Daisy created the correct candidates but then too strictly filtered them out while keeping unrelated hits.

However, O55:H7 contains other λ phages (Kyle et al., 2012), two of them (Cp7 and Cp9) have high similarity (up to 99% BLAST identity) to Sp15 (Stx1) at the *yehV* phage insertion site in *EHEC* O157:H7. When we look more closely at the six identified candidates, we observe that all of them have acceptor positions matching an alternative phage insertion site (see Table S1 in Kyle et al. (2012) for details on the phage insertion sites). Among these six candidates, there is the one true candidate regarding the donor positions (1,283,673-1,288,079). The acceptor coordinates (1,741,535-1,744,926) belong to the λ phage Cp7.

The candidate with the true donor positions encloses a region that is 2,876 bp larger than the actual Stx part. A BLAST search of this additional part yields hits on shiga toxin genes and ORFs, and Stx phage and prophage genes, as well as four further hits to *S. dysenteriae* Sd197 (CP000034.1) that match the donor regions of the remaining five candidates. These hits suggest a phage-origin of this additional 2,876 bp (1,285,203-1,288,079) as well as these donor regions. This is supported by the high percentage of donor region read pairs matching an entry in the bacteriophage database (up to 97%). The percentage of phage database hits of the one remaining candidate is also around 97%, suggesting another alternative phage-site in *S. dysenteriae* Sd197 as well. The donor positions of all of the filtered out candidates with matching acceptor positions also all fall within the phage-region ranging from position 1,288,585 to 1,329,490 (data not shown).

So the true challenge in this case study is the fact that the HGT was phage mediated, and that both acceptor and donor have several alternative and occupied phage insertion sites. According to Asadulghani et al. (2009), the same set of bacteriophages can also occupy different phage insertion sites between individuals of the same bacterial strain, making it possible that our candidate (1,741,535-1,744,926 to 1,283,673-1,288,079) actually is the true or most likely candidate in this case. Given the currently available information, we cannot verify that the six phage-related candidates are correct, but there is also sufficient evidence to consider them as such.

#### Assembly

SOAPdenovo2 assembled 100,601 contigs, 14,897 of them covered by one of the 93,905 scaffolds (N50 of 150). *E. coli* O157:H7 carries Stx1 and Stx2 at distinct locations. However, assembly examination with BLAST suggests that both toxins have been assembled on the same scaffold.

Both toxin HGT region lie within a phage so it is difficult to ascertain the specific positions of the genes carried by the phage. For Alien Hunter this is difficult because the tool already (correctly) recognizes the phage sequence itself as a GI. We therefore count *true region ƒound* as yes, if the candidate region is overlapping the phage carrying the toxins. Alien Hunter finds both Stx regions when applied to the HGT organism, but one is the region with the lowest rank (see Table2, (B) EHEC). The tool finds 382 candidate regions when applied to the assembly but non is matching the scaffold with the HGT region. With BWA-MEM, we find 177 candidate regions. One region is overlapping the true HGT region but without breakpoint precision on the donor. The region is reaching into the repetitive genome part following the shiga-toxin region in *S. dysenteriae* Sd197. The results acquired via BWA-MEM also support our hypothesis of an alternative phage insertion site: The acceptor region of this hit is 1,741,843-1,742,439. For all three tools, the true HGT candidate is integrated into a phage and, hence, we assign breakpoint distance zero (0/0).

## 5 Discussion

To the best of our knowledge, Daisy is the first approach that allows HGT detection directly from NGS data without requiring a *de novo* assembled genome. It rather relies on detecting HGT boundaries via a split-read mapping approach from SV detection methods. It uses the acceptor and donor genomes of the HGT as reference, and integrates coverage and read pair information for HGT candidate evaluation. Daisy facilitates applications related to outbreak scenarios of HGT related pathogens like, e.g., detection of novel bacterial strains evolved through HGT or the distinction of a single infection with such a strain from a parallel infection by two different strains.

We critically evaluated Daisy on three data sets. It has been often noticed that SV detection methods are hard to evaluate since the existence and exact positions of breakpoints are often not known. This is particularly true also for HGT events. In this study, we therefore focused on one simulated and two real data sets for which we also provide partial ground truth for future comparison. The data sets were chosen to both show the power of the approach but also to explore the limitations and provide guidance for other experiments. On the simulated *H. Pylori* data, Daisy produced the correct true positive candidate without false positives. For the real KO11FL data, the five single boundaries with the highest total split-read support already cover all five HGT related boundaries and have a distinctly higher support than the first false positive hit. For the real EHEC data, we called six candidates which all fall into alternative phage insertion sites in both acceptor and donor. The alternative assembly only produced meaningful assemblies for *H. pylori* and EHEC. On the *H. pylori* data set, Alien Hunter and BWA-MEM both found the HGT region as the best candidate, but Alien Hunter with low breakpoint precision and many alternative hits. On the EHEC data set, only BWA-MEM found the true candidate on the assembly data but with more FPs than Daisy. An approach of mapping based HGT detection integrating several SV detection methods is therefore a highly useful strategy. The EHEC example, where we reduced the 145 pure split-based candidates to a few candidates with required HGT signatures, shows how our candidate evaluation successfully filters out false positive hits. Although these use cases were overall successful, the results also show some challenges and need for future development, which we outline below.

One prerequisite of the current approach is acceptor and donor genomes that are involved in the HGT event are known. Selecting these candidate genomes given a set of reads from the HGT carrier genome is a challenging task of its own. It is closely related to the metagenomics problem of finding all occurring species contained in a sample given a set of reads, and is a crucial pre-step for applications in diagnostics. Thus, tools such as MicrobeGPS (Lindner and Renard, 2015) or Kraken (Wood and Salzberg, 2014) can serve to identify candidates for follow-up analysis with our tool.

In this first version of Daisy, we focused on the idea of using mapping-based evidence such as coverage and read pair signatures. Existing parametric HGT detection methods use genome signatures such as differing GC content (Daubin et al., 2003), atypical codon usage (Lawrence and Ochman, 2002) or *k-mer* frequencies (like Alien Hunter) for identification. In a more comprehensive future version, and in cases when the HGT organism reference genomes are available, these genome signatures are possible further filtering options of candidate regions.

Currently, automated filtering and HGT candidate evaluation is only available for a single donor genome. More complex, decomposite HGT regions, consisting of multiple genes from various donor genomes such as in the KO11FL example, require more sophisticated combination and evaluation of candidates and paired support across the donors. An automated extension could benefit the application also in the context of, e.g., genetically modified organisms. While the detection is possible with our approach, as seen in the KO11FL, more manual investigation is required.

It should be noted that Daisy relies on a mapping approach to known reference genomes. In recent HGT events or artificial gene transfers, the mapping to reference genomes is easier than for longer evolutionary time spans. As a result, HGT boundaries are more obvious and identifiable with higher confidence without strong influences of evolution. The example on the EHEC data shows that the ongoing fast evolution of bacteria makes HGT boundaries fuzzy, parallel HGT events obscure boundaries, and HGTs mediated by phages make the area around the target gene ambiguous and evaluation difficult. In general, our approach should be seen a step towards a more comprehensive analysis pipeline of sequencing data where multiple complementary methods are integrated. The goal of such a pipeline would be a full investigation of a complete bacterial genome with, e.g., genome annotation and classification, SNP and SV characterization, HGT detection and more.

## 6 Conclusion

With our tool Daisy, we present the first mapping-based HGT detection approach known so far. Our approach shows sound results with base pair precision for simulated and real data sets. Alternative assembly give supportive results but was not successful for all data sets. Daisy was built for and evaluated on bacteria, but should in principle also be applicable for HGT detection in other organisms such as plants. Daisy is written in Python as a ready to use tool building on NGS standard input and output formats and is freely available from http://github.com/ktrappe/daisy.

## Acknowledgement

We thank Knut Reinert and Martin Lindner for inspiring discussions, and Enrico Seiler and Jan R. Forster for their assistance in code development and innervating discussions.

## Author’s Contributions

KT participated in the design of the study, developed the pipeline, analyzed data sets and drafted the manuscript. TM jointly conceived of the study with BYR, participated in its design and helped writing the manuscript. BYR participated in the design of the study, pipeline development and manuscript editing. All authors read and approved the final manuscript.

Conflict of interest: none declared.

